# The relationship between brain atrophy and cognitive-behavioral symptoms in retired Canadian football players with multiple concussions

**DOI:** 10.1101/261404

**Authors:** Karen Misquitta, Mahsa Dadar, Apameh Tarazi, MW Hussain, MK Alatwi, Ahmed Ebraheem, Namita Multani, Mozhgan Khodadadi, Ruma Goswami, Richard Wennberg, Charles Tator, Robin Green, Brenda Colella, Karen Davis, David Mikulis, Mark Grinberg, Christine Sato, Ekaterina Rogaeva, D. Louis Collins, Maria Carmela Tartaglia

## Abstract

Multiple concussions, particularly in contact sports, have been associated with cognitive deficits, psychiatric impairment and neurodegenerative diseases like chronic traumatic encephalopathy. We used volumetric and deformation-based morphometric analyses to test the hypothesis that repeated concussions may be associated with smaller regional brain volumes, poorer cognitive performance and behavioural symptoms among former professional football players compared to healthy controls. This study included fifty-three retired Canadian Football League players, 25 age- and education-matched healthy controls, and controls from the Cambridge Centre for Aging and Neuroscience database for validation. Volumetric analyses revealed greater hippocampal atrophy than expected for age in former athletes with multiple concussions than controls and smaller left hippocampal volume was associated with poorer verbal memory performance. Deformation-based morphometric confirmed smaller bilateral hippocampal volume that were associated with poorer verbal memory performance in athletes. Repeated concussions may lead to greater regional atrophy than expected for age.

## Introduction

There is a high incidence of concussions, particularly among players of contact sports, with an estimated 1.6 to 3.8 million sports-related concussions occurring each year in the United States alone^1^. Professional players of contact sports will experience hits to the head but not all will report having a concussion. While most concussive events resolve within weeks, at least 10% of patients experience prolonged symptoms known as post-concussion syndrome^2^. Recently, there is growing concern that repeated concussions can cause late life mild cognitive impairment, an earlier onset of Alzheimer’s disease^3^ or the neurodegenerative disease called chronic traumatic encephalopathy (CTE)^4^. The majority of CTE cases have been reported in athletes involved in contact sports, including boxing, football, hockey, rugby, wrestling and soccer^4-6^.

Neuronal damage from traumatic brain injury (TBI) has been associated with cerebral atrophy in studies of mild, moderate and severe brain injury^7^. Normal aging is also associated with mild brain volume loss and some cognitive deficits^8^. Accelerated cognitive decline may occur as a result of mild, moderate or severe TBI, and exacerbate deficits associated with the normal aging process^9^. Memory impairment is one of the most frequent cognitive complaints following mild, moderate and severe TBI^10^. Verbal memory impairment may result from injury to the left medio-temporal and hippocampal regions^11^ while deficits in visuospatial memory may be associated with these regions in the right hemisphere^12^. Moreover, post-concussive symptoms include behaviour and personality changes, such as depression, apathy, impulsivity and aggression^13^, which have been associated with generalized and regional brain atrophy in various study populations^14^.

Several neuroimaging techniques have been used to examine whether symptoms resulting from multiple concussions are associated with detectable changes in brain volume and function^15-19^. However, results from these studies have been mixed; while some identify structural and functional brain changes associated with symptoms in both acute and chronic concussed populations^15,17,18^, other studies report no abnormalities^16,19^.

Severe, moderate and mild TBI are associated with long-term damage to the brain^20,21^. We hypothesize that long-term damage can result from mild TBI and contribute to measurable brain atrophy and that this atrophy will be associated with cognitive deficits and behavioural changes. Using structural segmentation and deformation-based morphometry (DBM)^22^ analyses, the current study compares the effect of multiple concussions on regional brain volumes in retired professional athletes from the Canadian Football League (ex-CFL) with non-athlete control subjects with no history of concussion. As the sample size of our control group was small, a larger control population from the Cambridge Centre for Aging and Neuroscience database was leveraged for further analysis. We also investigated the relationship between regional brain volume and memory and personality changes. We predict that ex-CFL will show greater focal atrophy, and these regions will be associated with poorer memory performance and personality changes compared to controls.

## Results

Subject demographics are listed in Table 1. There were no significant between-group differences in age, education, or memory score. There was also no difference in the proportion of *APOE*-e4 allele carriers between the groups. Number of years playing professional football was significantly related to smaller left and right hippocampus and left and right amygdala volumes, but not with ventricular volume (Table 2). After Bonferroni correction for multiple comparisons, only correlations with the left and right amygdala remained significant (p<0.01).

**Table 1.**
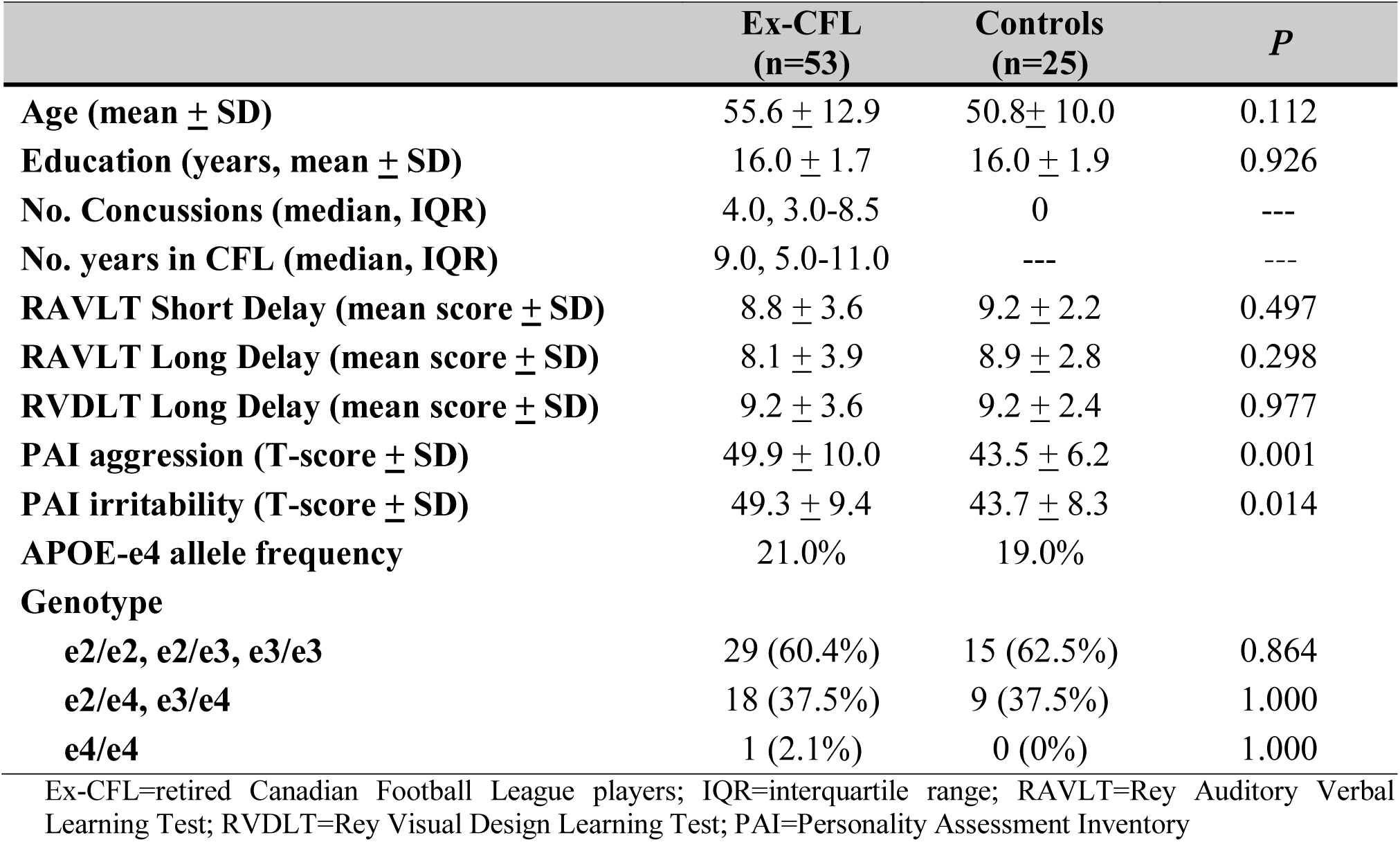
Subject demographics

**Table 2.** Correlation coefficients for years playing professional football in ex-CFL players.

| **Variables** | **Years played in CFL** |  |
| --- | --- | --- |
|  | **r** | **p** |
| **Left Hippocampus** | -0.321 | 0.022 |
| **Right Hippocampus** | -0.339 | 0.017 |
| **Left Amygdala** | -0.448 | 0.002 |
| **Right Amygdala** | -0.397 | 0.006 |
| **Ventricle** | 0.212 | 0.096 |

CFL=retired Canadian Football League

Linear regression was used to measure the relationship between age and right and left hippocampal volumes in ex-CFL, study controls and control participants from the Cam-CAN database. Different intercepts and interactions with age were allowed for each cohort. The linear regression model was:

H=α_0_+ α_1_ Cohort_study controls_ + α_2_Cohort_ex-CFL_ +β_0_Age+β_1_Age:Cohort_study controls_+ β_2_ Age:Cohort_ex-CFL_ Where “:” indicates an interaction between the two terms, H is the volume of the hippocampi (left or right hippocampus), α_0_ is the intercept term for Cam-CAN cohort, α_1_ is the additional intercept for the study controls cohort, and α_2_ is the additional intercept for the ex-CFL cohort. Similarly, β_0_ is the linear slope for age for Cam-CAN cohort, and β_1_ and β_2_ are the additional slopes for study controls and ex-CFL cohort, respectively. A negative β value indicates a relatively steeper slope (added to the negative slope for age estimated from the Cam-CAN cohort) for the respective cohort. Similarly, a positive β value indicates a less steep slope in comparison with Cam-CAN cohort. The estimated parameters of the model are reported in Table 4.

Linear regression showed a statistically significant association between age and the volumes of both the left hippocampus and the right hippocampus in Cam-CAN controls. This association was significantly different (i.e. steeper slope for both the left hippocampus and the right hippocampus) in the ex-CFL players, but not for study controls (Table 3; Figure 1).

**Figure 1.**
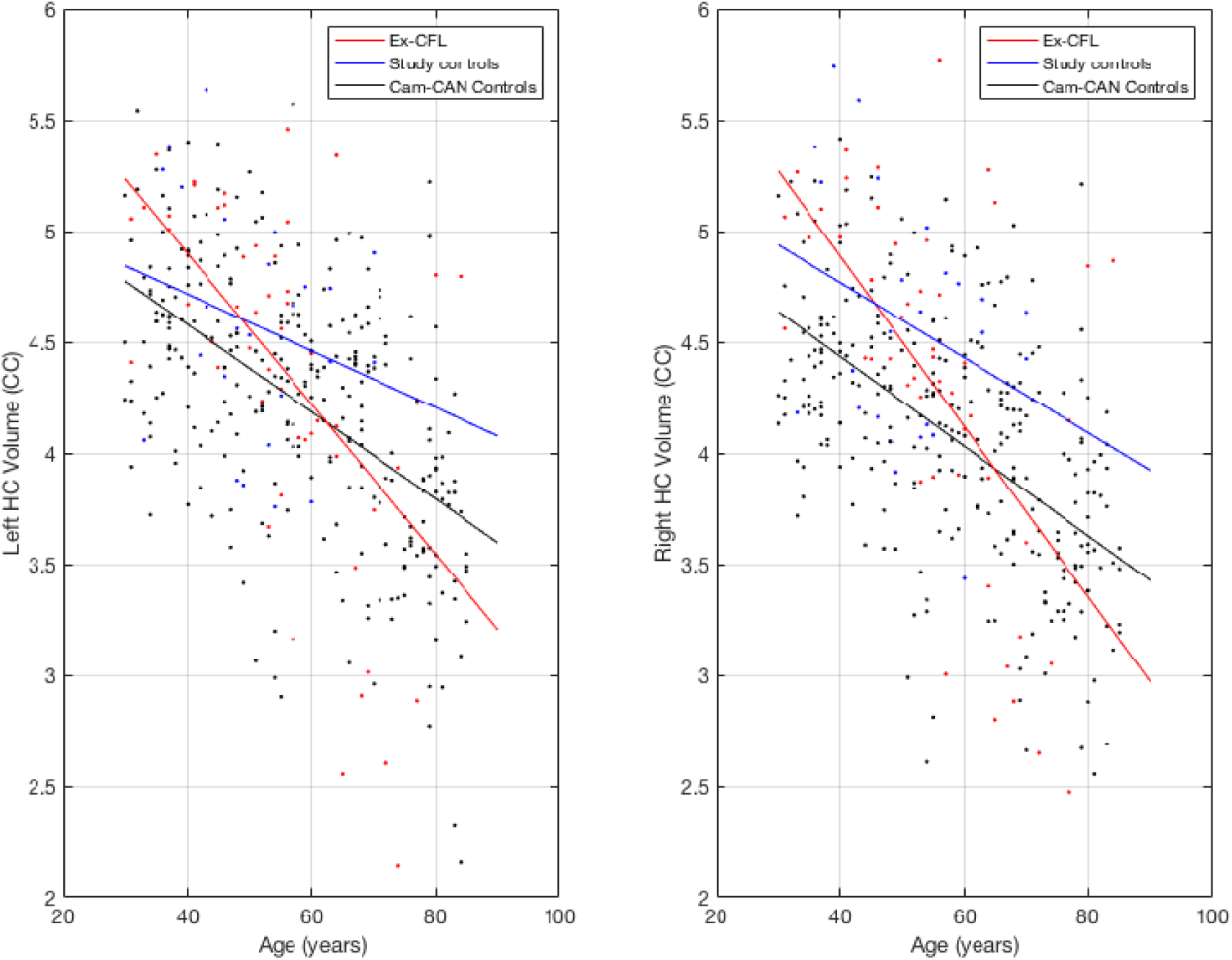
Linear regression model results showing the relationship between left and right hippocampal volumes and age in ex-CFL players, study controls and Cam-CAN controls. Modelling of age and left and right hippocampal volumes show a much steeper effect of age on hippocampal volumes in the former CFL players compared to both study controls and Cam-CAN controls.

**Table 3.** Linear regression for left and right hippocampal volume with age in ex-CFL players, study controls and Cam-CAN controls, where “:” indicates an interaction between the two terms.

| Variable | Left Hippocampus |  | Right Hippocampus |  |
| --- | --- | --- | --- | --- |
| | $\beta$ | $p$ | $\beta$ | $p$ |
| **Intercept**<sub>Cam-CAN controls</sub> | -0.046 | 0.350 | -0.077 | 0.112 |
| **Cohort**<sub>study controls</sub> | 0.412 | 0.047 | 0.603 | 0.003 |
| **Cohort**<sub>ex-CFL players</sub> | 0.119 | 0.347 | 0.213 | 0.084 |
| **Age**<sub>Cam-CAN controls</sub> | -0.484 | <0.001 | -0.479 | <0.001 |
| **Age: Cohort**<sub>study controls</sub> | 0.169 | 0.528 | 0.078 | 0.765 |
| **Age: Cohort**<sub>ex-CFL players</sub> | -0.351 | 0.017 | -0.431 | 0.003 |

Ex-CFL=retired Canadian Football League players

To assess whether hippocampal volume was associated with memory function, we investigated the relationship between left hippocampal volume and performance on the RAVLT (verbal memory task) and right hippocampal volume and RVDLT (visual memory task). The ex-CFL group, but not the study control group, showed a significant relationship between smaller left hippocampal volume and poorer word recall performance on the RAVLT short delay (r=0.523, p=0.001) and RAVLT long delay scores (r=0.447, p=0.002) (Figure 2A and 2B). These correlations remained significant after Bonferroni correction (p<0.016). There were no significant correlations between RVDLT long delay scores and right hippocampal volume in either the ex-CFL or control groups (Figure 2C). There were no significant differences between ex-CFL players and normal controls on RAVLT short delay scores (*p*=0.497) long delay scores (*p*=0.298), or RVDLT long delay scores (*p*=0.977) (Table 1).

**Figure 2.**
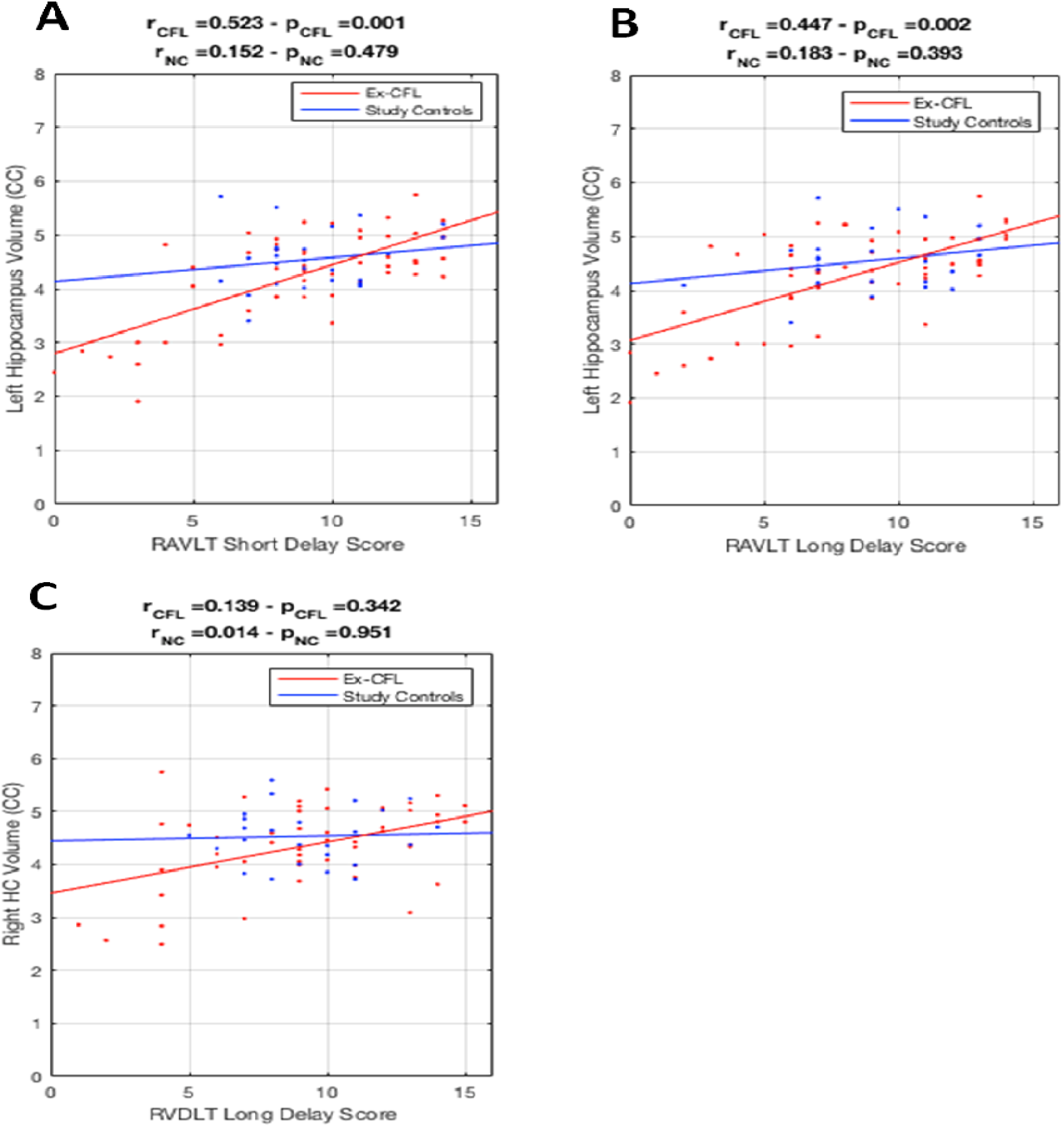
Pearson correlation graphs in ex-CFL and controls between left hippocampal volumes and A) the RAVLT short delay total score, and B) the RAVLT long delay total score, and between right hippocampal volume and C) the RVDLT long delay total score. Significant relationships were found between left and right hippocampal volumes and RAVLT short and long delay scores in the ex-CFL but not in the study controls.

DBM analysis was used to identify which areas across the brain may be associated with memory performance. Correlations with DBM maps also showed a significant relationship with RAVLT short delay in the ex-CFL players (Figure 3), where poorer scores on recall after a short delay were associated with smaller left and right hippocampal regions, after FDR correction (r=0.552, p=0.026; r=0.492, p=0.041, respectively).

**Figure 3.**
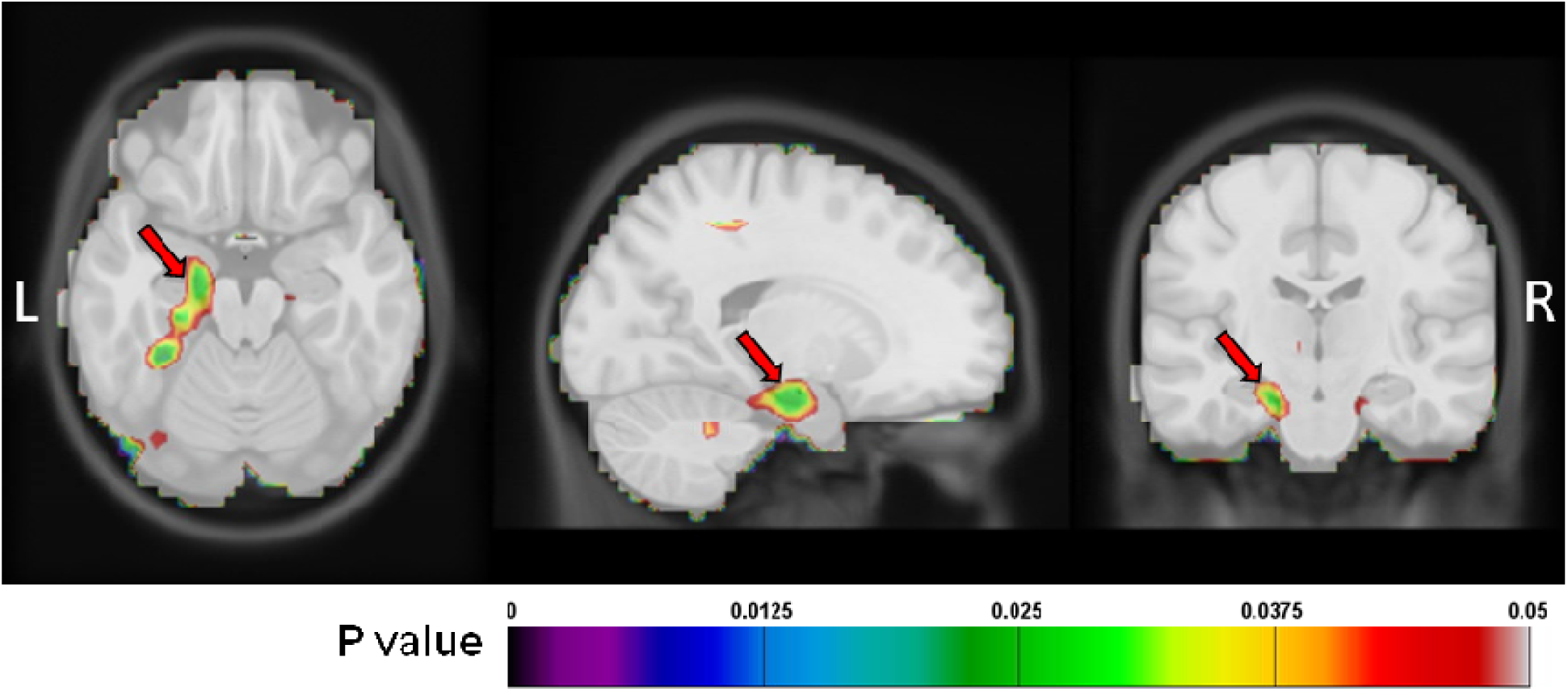
Deformation-based morphometry showing regions associated with RAVLT short delay scores in ex-CFL players. FDR (q=0.05) corrected p value map for regions associated with RAVLT short delay scores in ex-CFL players. Colours represent p-values. Red arrows indicate the left medial temporal lobe, including primarily the hippocampus, parahippocampus, and entorhinal cortex.

The relationship between amygdala volume and aggression and irritability personality traits from the PAI were also examined in ex-CFL players. Left and right amygdala volumes were not significantly associated with T-scores for aggression (*r*=0.162, *p*=0.255; *r*=0.206, *p*=0.147, respectively) and irritability (*r*=0.204, *p*=0.150; *r*=0.192, *p*=0.177, respectively) measured by the PAI. Ex-CFL showed higher T-scores on the PAI for aggression (p=0.001) and irritability (p=0.014) in this study compared to control subjects.

## Discussion

Using DBM and volumetric analyses, this study examined the effect of repeated concussions on regional brain volumes, cognitive performance and behavioural symptoms among former professional football players compared to healthy controls. Volumetric analysis demonstrated a significantly greater effect of age on brain volume reduction among retired professional football players compared with healthy controls. More specifically, the effect of age on hippocampal volume reduction was significantly greater in the ex-CFL cohort compared to the study controls and Cam-CAN control subjects. Our findings confirm an effect of age on brain volume, and this effect appears to be amplified in the ex-CFL group. Moreover, smaller left hippocampal volume was associated with worse performance on the short and long delay verbal memory scores of the RAVLT. In the ex-CFL players, DBM analysis also showed smaller bilateral hippocampal volume associated with poorer verbal memory performance. Although ex-CFL scored significantly higher than controls for symptoms of irritability and aggression on the PAI compared to controls, they were not in an elevated range^23^. We did not find any relationship between PAI irritability and aggression scores and amygdala volume in our ex-CFL or control groups.

Number of career years playing football with the CFL was associated with smaller hippocampi and amygdala volumes and larger ventricular volume in the ex-CFL. Career years were examined as a proxy for number of concussions because concussion history was self-reported by athletes and therefore a less reliable measure as a result of recall bias. Greater years playing professional football increases exposure to concussions and may contribute to the effect of age on brain volume observed in our ex-CFL group. In addition, many concussions may go unrecognized by athletes and/or their coaches as symptoms generally resolve on their own^24^. In our study, the four players reporting no concussions each played between 9-12 years in the CFL. It is possible that professional players of contact sports will experience hits to the head but may not report having a concussion, especially if they do not experience persistent post-concussive symptoms. As well, since concussion can be associated with both anterograde and retrograde amnesia, the players may have forgotten them^25^.

Previous studies have found evidence of cerebral atrophy in individuals with mild,moderate and severe TBI^26,27^. In particular, ventricular enlargement and volume loss of the hippocampi and thalami have been reported in studies of mild, moderate and severe TBI^8,28^. Due to its location in the middle cranial fossa, the hippocampus is situated in a region vulnerable to injury^7^, and hippocampal atrophy following mild, moderate and severe TBI has been reported in both animal^29^ and human studies. The current study suggests that the effect of age on hippocampal volume loss is greater in those who have experienced repeated concussions. Moreover, hippocampal atrophy is associated with memory impairment in normal aging as well as in various neurodegenerative diseases including Alzheimer’s disease and frontotemporal lobar degeneration. As would be expected, we found a relationship between smaller bilateral hippocampal volume and poorer performance on both early and delayed recall of a verbal memory task in ex-CFL players, and this relationship was greater in ex-CFL compared to study controls.

APOE protein is involved in lipid metabolism; and the *APOE*-e4 allele increases risk of Alzheimer’s disease in a dose-dependent fashion (by three times for heterozygotes), whereas the *APOE*-e2 allele confers a protective benefit^30^. Some studies have suggested that *APOE-*e4 allele may be also associated with less efficient cognitive processing in concussed athletes^31^. However, in the current study, we did not find a significant difference between ex-CFL and control groups in *APOE* genotype or allele frequency.

Irritability and aggression are often reported in cases of CTE^20^, and their relationship to amygdala volume was examined in this study. Significantly higher T-scores for both symptoms were found in ex-CFL compared to controls, but they were within the normal range in both groups. We did not find a significant association between amygdala volume and PAI T-scores for aggression and irritability.

At present, few studies have looked at changes in regional cerebral atrophy among retired athletes with a history of multiple concussions. Results from these studies have been mixed, with some detecting no significant abnormalities in concussed athletes^16^. Most studies also include participants with a range of mild to severe TBI with few studies having looked at structural brain changes in concussion alone. Here, we show differences in brain volume in a group of athletes who were exposed to repetitive head impacts and a history of multiple concussions only.

The current study uses DBM to explore structural brain changes that may result from repeated exposure to concussive head injury. DBM is a whole brain analysis that allows the examination of macroscopic differences across the entire brain by comparing the position of each voxel to a standard brain and so may be more sensitive to subtle volume changes^22,32^. Studies have examined the trajectory of grey matter atrophy with age using both linear and nonlinear models with mixed results and variation among structures^33^. Further work is needed to better understand the effect of age on atrophy in healthy and concussed populations.

Our study is limited in its ability to detect causal and temporal relationships in regional cerebral atrophy over time due to its cross-sectional design. Future studies should include longitudinal designs that can better assess causal relationships. In addition, the number of concussions was self-reported by ex-CFL players and these numbers are therefore subject to response and recall bias. Ex-CFL players were also self-referred and it is possible that those with specific neurobehavioural symptoms are more likely to choose to participate in this study. Participants in the current study were restricted to male professional football players and our findings may not apply to other groups including females and non-athletes. Age since retirement may also be a confounder in our analysis, however it would be difficult to de-correlate with age. Other confounders including family history of dementia, alcohol and substance abuse, and use of performance enhancing drugs were not included in our analysis but may be important modulators of the age-related changes in volume observed in this study, and future work should assess the potential contribution of these confounders.

We have demonstrated greater age effects on hippocampal volume in ex-CFL players. Moreover, we showed that these changes are associated with the number of career years playing for the CFL. Taken together, these findings suggest that multiple concussions may contribute to pathological changes that are associated with greater age-related atrophy and result in earlier focal atrophy, although longitudinal studies are needed to determine the essence of this relationship. Future studies that track structural, cognitive and behavioural changes over time can provide additional insight into the effects of concussions on brain structure and function.

## Materials and methods

### Participants

This study included 53 ex-CFL players (mean age=55.6±12.9 years), most of whom report multiple concussions, and 25 healthy age- and education-matched male non-concussed controls (mean age=50.8±10.0 years), recruited from the general population. Athletes played for one or more seasons with the CFL. Informed consent was obtained and the study was approved by the University Health Network research ethics board.

Age-matched male controls from the Cambridge Centre for Aging and Neuroscience (Cam-CAN, N=321, mean age of 58.1±16.0 years, range 30-85) were used for validation due to the small size of the local healthy control group. Data were obtained from the Cam-CAN repository (available at http://www.mrc-cbu.cam.ac.uk/datasets/camcan/)^34,35^.

The median number of self-reported concussions in the ex-CFL group was 4 (Table 1). Exclusion criteria included: neurological disorders prior to concussions (e.g. seizure disorder), systemic illnesses known to affect the brain (e.g. diabetes and lupus), a history of psychotic disorder, known developmental disorders, and history of migraines. Similar criteria were used for study and Cam-CAN controls^35^. Concussion exposure was based on players’ recall of injury during a semi-structured interview in accordance with the Zurich Guidelines on Concussions^36^. Absence of concussions in the control group was verified through interview with control subjects.

### Neuropsychological Assessment

All participants underwent an extensive neuropsychological test battery comprising a series of cognitive and behavioural assessments. Memory was assessed by the Rey Auditory Verbal Learning Test (RAVLT)^37^, which is a test of verbal learning and memory; and by the Rey Visual Design Learning Test (RVDLT)^38^ assessing visual learning and memory. For the RAVLT, participants were asked to repeat 15 unrelated words over five consecutive trials, after which an interference list is presented and recalled. Subjects are then asked to recall the original list after this short delay, and again after a 20-minute long delay. The number of words recalled after the short and after the long delay were the primary behavioural outcome measures. For the RVDLT, participants were presented with 15 stimulus cards with geometric design (over five consecutive trials), and asked to draw all designs they could recall after each trial. Twenty minutes after completing the final trial, participants were asked to redraw as many of the 15 designs they could recall. The number of accurately drawn figures was the primary behavioural outcome measure.

Symptoms measured by the Personality Assessment Inventory (PAI)^23^ were included for analysis, correcting for age. The PAI was chosen to measure personality changes frequently associated with concussion. Aggression and irritability were chosen for analysis as the PAI symptoms most relevant to concussion. The PAI is a comprehensive and informative self-report questionnaire of adult personality and psychopathology, and contains 344 items scored on a 4-point scale: F=false, ST=slightly true; MT=mainly true; VT=very true. This assessment contains 22 full scales (four validity scales, 11 clinical scales, five treatment scales, and two interpersonal scales) with 10 of these scales further subdivided into 31 conceptually derived subscales. *T*-scores are based on a census matched standardization sample of 1,000 normal adults.

### Neuroimaging

#### IMAGE ACQUISITION

Participants underwent a whole-brain scan using a T1-weighted inversion recovery prepped, 3-dimensional IR-FSPGR (inversion fast spoiled gradient echo) sequence at 3 Tesla (GE Signa HDx, Milwaukee, WI, USA) with the following parameters: 180 axial slices, 1x1x1-mm voxels, 256x256 matrix size, 25.6-cm field of view, flip angle=158°, echo time=3ms, repetition time=7.8ms, inversion time = 450ms.

Cam-CAN participants underwent T1-weighted MPRAGE (magnetization prepared rapid acquisition gradient echo) sequence at 3 Tesla (Siemens TIM Trio scanner with a 32-channel head coil) with the following acquisition parameters: 1x1x1-mm voxels, field of view=256x240x192, flip angle=9°, echo time=2.99ms, repetition time=2250ms, inversion time=900ms.

#### PRE-PROCESSING

T1-weighted scans of the subjects were pre-processed through our standard pipeline. Image denoising^39^, intensity non-uniformity correction^40^, and image intensity normalization into range (0-100) using histogram matching were performed.

#### DEFORMATION-BASED MORPHOMETRY

DBM analysis was performed using MNI MINC tools. Pre-processed images were first linearly (using a 9-parameter rigid registration)^41^ and then non-linearly warped^42^ to an average template brain of 152 healthy young individuals (MNI-ICBM-152). The local deformation obtained from the non-linear transformations was used as a measure of tissue expansion or atrophy. DBM was used to examine the relationship between brain volume and performance on a memory task. Voxel-wise deformation maps were correlated with RAVLT short delay scores and corrected for multiple comparisons using False Discovery Rate (FDR), thresholded at q=0.05.

#### ANALYSIS OF SUBCORTICAL STRUCTURES

All images were first linearly (using a 9-parameter rigid registration) and then nonlinearly registered to an average template (MNI ICBM152) as part of the ANIMAL software^43,44^. The deep structures, i.e., thalami, ventricles, putamen, and caudate, were segmented as part of the validated ANIMAL software by warping segmentations from ICBM152 back to each subject using the obtained nonlinear transformations. The hippocampi and amygdala were segmented using a validated automated patch-based label-fusion technique^44^. The method selects the most similar templates from a library of labelled MRI template images, and combines them with majority voting scheme to assign the highest weighted label to a given voxel to generate a discrete segmentation. Quality control was performed on the individual registered images as well as the automated structure segmentations by visual inspection. The volumes of the structures were then calculated from the segmentations in the ICBM152 space, i.e. the values were scaled by a scaling factor inversely proportional to the intracranial volume to account for differences in head sizes.

### *APOE*genotyping

For all study participants, genomic DNA was extracted from whole blood using Qiagen kits. The two single nucleotide polymorphisms in *APOE* (rs7412 and rs429358) defining the *APOE*-e2, -e3 and -e4 alleles were genotyped as previously described^29^.

### Statistical Analysis

Statistical analyses were performed using MATLAB R2015b software (MATLAB, Natick, MA, USA). The relationship between hippocampal volume and age was examined by multiple linear regression. Partial correlations (two-tailed) were calculated between hippocampal, amygdala and ventricular volumes and years playing football, correcting for age, and between these volumes and RAVLT short and long delay scores and RVDLT long delay scores, correcting for age and education. A Student’s t-test for independent samples was used to compare age, education, RAVLT and RVDLT scores, and PAI sores for aggression and irritability symptoms between the ex-CFL players and the control group. Chi-square or Fisher’s exact test were used to compare APOE genotype, between ex-CFL players and controls. Pearson correlations (two-tailed) were also calculated between behavioural symptoms (aggression and irritability) and amygdala volume, correcting for age. All images were linearly transformed into the same space before analysis, thus accounting for head size. The significance level for all analyses was set at p<0.05 (two-tailed). Bonferonni correction for multiple comparisons was performed.

## Acknowledgements

This study was funded by the Physicians’ Services Incorporated Foundation and the Toronto General and Western Hospital Foundation through specific donations to the Canadian Concussion Centre. Data collection and sharing for this project was provided by the Cambridge Centre for Ageing and Neuroscience (CamCAN). CamCAN funding was provided by the UK Biotechnology and Biological Sciences Research Council (grant number BB/H008217/1), together with support from the UK Medical Research Council and University of Cambridge, UK. KM funded by CIHR. DLC received a grant from the André Carron Family. DLC and MCT contributed to study concept and design. KM, MD, AT, MWH, MKA, AE, NM, MK, R Goswami, RW, CT, R Green, BC, KD, MG, CS, ER, DM, and MCT contributed to data acquisition and analysis. KM, MD and MCT contributed to preparing the manuscript. We thank all the participants for the generous contribution of their time.

## References

1. Langlois JA, Rutland-Brown W, Wald MM (1991) The epidemiology and impact of traumatic brain injury: a brief overview. J Head Trauma Rehabil 21(5):375–8.

2. Hiploylee C, Dufort PA, Davis HS, Wennberg RA, Tartaglia MC, Mikulis D, Hazrati LN, Tator CH (2017) Longitudinal Study of Postconcussion Syndrome: Not Everyone Recovers. J Neurotrauma 34(8):1511–23. doi:10.1089/neu.2016.4677

3. Abner EL, Nelson PT, Schmitt FA, Browning SR, Fardo DW, Wan L, Jicha GA, Cooper GE, Smith CD, Caban-Holt AM, Van Eldik LJ, Kryscio RJ (2014) Self-reported head injury and risk of late-life impairment and AD pathology in an AD center cohort. Dement Geriatr Cogn Disord 37(5-6):294–306. doi:10.1159/000355478

4. Omalu BI, DeKosky ST, Minster RL, Kamboh MI, Hamilton RL, Wecht CH (2005) Chronic traumatic encephalopathy in a National Football League player. Neurosurgery 57(1):128–34.

5. Omalu BI, DeKosky ST, Hamilton RL, Minster RL, Kamboh MI, Shakir AM, Wecht CH (2006) Chronic traumatic encephalopathy in a national football league player: part II. Neurosurgery 59(5):1086–92; discussion 92-3.

6. Omalu BI, Fitzsimmons RP, Hammers J, Bailes J (2010) Chronic traumatic encephalopathy in a professional American wrestler. J Forensic Nurs 6(3):130–6. doi:10.1111/j.1939-3938.2010.01078.x.

7. Shenton ME, Hamoda HM, Schneiderman JS, Bouix S, Pasternak O, Rathi Y, Vu MA, Purohit MP, Helmer K, Koerte I, Lin AP, Westin CF, Kikinis R, Kubicki M, Stern RA, Zafonte R (2012) A review of magnetic resonance imaging and diffusion tensor imaging findings in mild traumatic brain injury. Brain Imaging Behav 6(2):137–92. doi:10.1007/s11682-012-9156-5.

8. Bigler ED, Blatter DD, Anderson CV, Johnson SC, Gale SD, Hopkins RO, Burnett B (1997) Hippocampal volume in normal aging and traumatic brain injury. AJNR Am J Neuroradiol 18(1):11–23.

9. Tremblay S, De Beaumont L, Henry LC, Boulanger Y, Evans AC, Bourgouin P, Poirier J, Theoret H, Lassonde M (2013) Sports concussions and aging: a neuroimaging investigation. Cereb Cortex 23(5):1159–66. doi:10.1093/cercor/bhs102.

10. Rabinowitz, AR, Levin HS (2014) Cognitive Sequelae of Traumatic Brain Injury. Psychiatr Clin North Am 37(1): 1–11. doi:10.1016/j.psc.2013.11.004

11. Frisk V, Milner B (1990) The role of the left hippocampal region in the acquisition and retention of story content. Neuropsychologia 28: 349–359.

12. Smith ML, Milner B (1981) The role of the right hippocampus in the recall of spatial location. Neuropsychologia 19: 781–793.

13. Malia K, Powell G, Torode S (1995) Personality and psychosocial function after brain injury. Brain Inj 9(7):697–712.

14. Matthies S, Rusch N, Weber M, Lieb K, Philipsen A, Tuescher O, Ebert D, Hennig J, van Elst LT (2012) Small amygdala-high aggression? The role of the amygdala in modulating aggression in healthy subjects. World J Biol Psychiatry 13(1):75–81. doi:10.3109/15622975.2010.541282.

15. Goswami R, Dufort P, Tartaglia MC, Green RE, Crawley A, Tator CH, Wennberg R, Mikulis DJ, Keightley M, Davis KD (2016) Frontotemporal correlates of impulsivity and machine learning in retired professional athletes with a history of multiple concussions. Brain Struct Funct 221:1911–1925. doi:10.1007/s00429-015-1012-0.

16. Ilvesmaki T, Luoto TM, Hakulinen U, Brander A, Ryymin P, Eskola H, Iverson GL, Ohman J (2014) Acute mild traumatic brain injury is not associated with white matter change on diffusion tensor imaging. Brain 137(Pt 7):1876–82. doi:10.1093/brain/awu095.

17. Meier TB, Bergamino M, Bellgowan PS, Teague TK, Ling JM, Jeromin A, Mayer AR (2016) Longitudinal assessment of white matter abnormalities following sports-related concussion. Hum Brain Mapp 37(2):833–45. doi:10.1002/hbm.23072.

18. Multani N, Goswami R, Khodadadi M, Ebraheem A, Davis KD, Tator CH, Wennberg R, Mikulis DJ, Ezerins L, Tartaglia MC (2016) The association between white-matter tract abnormalities, and neuropsychiatric and cognitive symptoms in retired professional football players with multiple concussions. J Neurol 263(7):1332–41. doi:10.1007/s00415-016-8141-0.

19. Tremblay S, Beaule V, Proulx S, Tremblay S, Marjanska M, Doyon J, Lassonde M, Theoret H (2014) Multimodal assessment of primary motor cortex integrity following sport concussion in asymptomatic athletes. Clin Neurophysiol 125(7):1371–9. doi:10.1016/j.clinph.2013.11.040.

20. McKee AC, Cantu RC, Nowinski CJ, Hedley-Whyte T, Gavett BE, Budson AE, Santini VE, Lee HS, Kubilus CA, Stern RA (2009) Chronic Traumatic Encephalopathy in Athletes: Progressive Tauopathy following Repetitive Head Injury. J Neuropathol Exp Neurol 68(7): 709–735. doi:10.1097/NEN.0b013e3181a9d503.

21. Corsellis, JA, Bruton CJ, Freeman-Browne D (1973) The aftermath of boxing. Psychol Med 3: 270–303.

22. Ashburner J, Hutton C, Frackowiak R, Johnsrude I, Price C, Friston K (1998) Identifying global anatomical differences: deformation-based morphometry. Hum Brain Mapp 6(5-6):348–57.

23. Morey LC (1991) Personality Assessment Inventory professional manual. Psychological Assessment Resources.

24. Daneshvar DH, Nowinski CJ, McKee A, Cantu RC (2011) The Epidemiology of Sport-Related Concussion. Clin Sports Med 30(1):1–17. doi:10.1016/j.csm.2010.08.006.

25. Cantu RC (2001) Posttraumatic retrograde and anterograde amnesia: pathophysiology and implications in grading and safe return to play. J Athl Train 36(3):244–248.

26. Levine B, Kovacevic N, Nica EI, Cheung G, Gao F, Schwartz ML, Black SE (2008) The Toronto traumatic brain injury study: injury severity and quantified MRI. Neurology 70(10):771–8. doi:10.1212/01.wnl.0000304108.32283.aa.

27. MacKenzie JD, Siddiqi F, Babb JS, Bagley LJ, Mannon LJ, Sinson GP, Grossman RI (2002) Brain atrophy in mild or moderate traumatic brain injury: a longitudinal quantitative analysis. AJNR Am J Neuroradiol 23(9):1509–15.23.

28. Wilde EA, Bigler ED, Pedroza C, Ryser DK (2006) Post-traumatic amnesia predicts long-term cerebral atrophy in traumatic brain injury. Brain Inj 20(7):695–9.

29. Hicks RR, Smith DH, Lowenstein DH, Saint Marie R, McIntosh TK (1993) Mild experimental brain injury in the rat induces cognitive deficits associated with regional neuronal loss in the hippocampus. J Neurotrauma 10(4):405–14.

30. Saunders AM, Strittmatter WJ, Schmechel D, George-Hyslop PH, Pericak-Vance MA, Joo SH, Rosi BL, Gusella JF, Crapper-MacLachlan DR, Alberts MJ et al. (1993) Association of apolipoprotein E allele epsilon 4 with late-onset familial and sporadic Alzheimer’s disease. Neurology 43(8):1467–72.

31. Merritt VC, Rabinowitz AR, Arnett PA (2017) The influence of the Apolipoprotein E (APOE) gene on subacute post-concussion neurocognitive performance in college athletes. Arch Clin Neuropsychol 24:1–11. doi:10.1093/arclin/acx051.

32. Gaser C, Nenadic I, Buchsbaum BR, Hazlett EA, Buchsbaum MS (2001) Deformation-based morphometry and its relation to conventional volumetry of brain lateral ventricles in MRI. Neuroimage 13(6 Pt 1):1140–5.

33. Fjell AM, Westlye LT, Grydeland H, Amlien I, Espeseth T, Reinvang I, Raz N, Holland D, Dale AM, Wahlhovd KB, Alzheimer Disease Neuroimaging Initiative (2013) Critical ages in the life course of the adult brain: nonlinear subcortical aging. Neurobiol Aging 34(10):2239–47. doi:10.1016/j.neurobiolaging.2013.04.006.

34. Shafto MA, Tyler LK, Dixon M, Taylor JR, Rowe JB, Cusack R, Calder AJ, Marslen-Wilson WD, Duncan J, Dalgleish T et al. (2014) The Cambridge Centre for Ageing and Neuroscience (Cam-CAN) study protocol: a cross-sectional, lifespan, multidisciplinary examination of healthy cognitive ageing. BMC Neurol 14(14):204. doi:10.1186/s12883-014-0204-1.

35. Taylor JR, Williams N, Cusack R, Auer T, Shafto MA, Dixon M, Tyler LK, Cam-Can, Henson RN (2017) The Cambridge Centre for Ageing and Neuroscience (Cam-CAN) data repository: Structural and functional MRI, MEG, and cognitive data from a cross-sectional adult lifespan sample. Neuroimage 144(Pt B):262–9. doi:10.1016/j.neuroimage.2015.09.018.

36. McCrory P, Meeuwisse W, Aubry M, Cantu B, Dvorak J, Echemendia R, Engebretsen L, Johnston K, Kutcher J, Raftery M et al. (2013) Consensus statement on concussion in sport: the 4th International Conference on Concussion in Sport held in Zurich, November 2013. Br J Sports Med 47(5):250–8. doi:10.1136/bjsports-2013-092313.

37. Rey A (1964) L’examen Clinique en Psychologie. Paris: Presses Universitaires de France.

38. Spreen O, Strauss E (1991) A compendium of neuropsychological tests: Administration, norms and commentary. New York, Oxford: Oxford University Press.

39. Coupe P, Yger P, Prima S, Hellier P, Kervrann C, Barillot C (2008) An optimized blockwise nonlocal means denoising filter for 3-D magnetic resonance images. IEEE Trans Med Imaging 27(4):425–41. doi:10.1109/TMI.2007.906087

40. Sled JG, Zijdenbos AP, Evans AC (1998) A nonparametric method for automatic correction of intensity nonuniformity in MRI data. IEEE Trans Med Imaging 17(1):87–97.

41. Collins DL, Neelin P, Peters TM, Evans AC (1994) Automatic 3D intersubject registration of MR volumetric data in standardized Talairach space. J Comput Assist Tomogr 18(2):192–205.

42. Collins DL, Evans AC, Holmes C, Peters TM (1995) Automatic 3D segmentation of neuro-anatomical structures from MRI. Comp Imag Vis 3:139–52.

43. Collins DL, Evans AC (1997) Animal: Validation and Applications of Nonlinear Registration-Based Segmentation. Int J Patt Recogn Artif Intell 11(1271).

44. Collins DL, Pruessner JC (2010) Towards accurate, automatic segmentation of the hippocampus and amygdala from MRI by augmenting ANIMAL with a template library and label fusion. Neuroimage 52(4):1355–66. doi:10.1016/j.neuroimage.2010.04.193.

